# Octopus “Hypnosis”: Inducing Tonic Immobility for Studying Local Sensorimotor Responses and Arm-Sucker Coordination

**DOI:** 10.1101/2024.04.22.590669

**Authors:** Ekaterina D. Gribkova, Jilai Cui, Rhanor Gillette

## Abstract

Effective methods of anesthesia for octopuses are essential to physiological studies as well as for their welfare in scientific research. However, commonly used forms of general anesthesia using ethanol, magnesium chloride, and similar agents have certain drawbacks. While these methods effectively induce still states in the octopus, they also affect the peripheral body and nervous system and are therefore less than optimal for studying local behavior in octopus arms and suckers. Further, stupefying effects outlast the anesthetized state. We explore an old, rarely used method of octopus “hypnosis” in which tonic immobility is induced as a complementary and sometime alternative method to general anesthesia, as well as being particularly suited to studies of local arm-sucker coordination. We modify the procedure for better handling, unimpeded respiration, and isolation of arm peripheral nervous system from the central nervous system (CNS). In the still state, an arm can be neurophysiologically isolated from the CNS by local Mg2+ injection, removing need for isolation by amputation. Exemplary studies of arm-sucker coordination and electrode placements are presented. Additionally, an intriguing phenomenon is observed where the induction of tonic immobility is notably diminished in cases of senescence. This modified procedure offers new convenience and directions for octopus neurobehavioral research.

## INTRODUCTION

Effective methods of anesthesia are important to physiological studies of octopus in scientific research as well as to their welfare. Anesthesia, defined as loss of sensation, is characterized by lack of response to noxious stimuli as well as a state of neural depression (Abbo et al., 2021; Grimm et al., 2015). However, while general anesthesia effectively induces still states in the octopus, it also affects the peripheral nervous system (PNS), markedly reducing or eliminating local reactions in the peripheral body, thereby making studies of local behavior, such as in arms and suckers, infeasible. Here we reproduce “octopus hypnosis”, or tonic immobility (TI), as a still reaction induced by mild restraint in a mouth-side up position, where local responses are preserved, and with significantly reduced stress and rapid recovery. Notably, TI may also decrease communication between peripheral and central nervous systems (CNS) of the octopus, thereby allowing minor invasive manipulations like local anesthesia, electrode implants, and prolonged measures of peripheral behavior in the arms and suckers.

### General Anesthesia

Ethanol in sea water at 0.5% - 2% is commonly used general anesthesia for octopuses (Butler-Struben et al., 2018). However, potential drawbacks of ethanol anesthesia have been observed. In particular, ethanol anesthesia sometimes causes obvious stress. Even with slow introduction of the ethanol solution, animals may become noticeably agitated (increased ventilation, jetting water, and escape behavior)(Andrews & Tansey, 1981; Winlow et al., 2018), and as anesthesia takes effect the octopuses may ink. We have also noticed a recurring problem with experiments during ethanol anesthesia: in several instances, an arm implanted with electrodes or isolated by amputation seems to show decreased spontaneous behavior and sensitivity after recent recovery from anesthesia. The arms of an octopus implanted with electrodes under ethanol anesthesia also appear to show injury responses, along with less fine coordination and significantly diminished muscular responses to arm stimuli, increased curling at arm tips, and tucking of the implanted arm beneath its web. This apparent increased injury response could be due to activation of sensitized nociceptive afferents in the arm following ethanol washout (Butler-Struben et al., 2018; Crook et al., 2013; Perez et al., 2017). Further, arms amputated under ethanol anesthesia and moved to fresh seawater show severely diminished responses to sensory stimuli and even electrical stimulation, possibly due to summating effects of ethanol toxicity and amputation trauma.

Magnesium chloride solution (MgCl_2_; 330 mM at pH 8, closely iso-osmotic to sea water and the animals’ blood,) is also commonly used for octopus general anesthesia and seems to lack some of the negative effects of ethanol anesthesia, such as active behavioral stress responses. However, the divalent magnesium ion is sticky and typically takes longer to wash out from exposed tissues than ethanol, so recovery from general MgCl_2_ anesthesia is quite slower. Moreover, both MgCl_2_ and ethanol anesthesia risk stopping the animal’s breathing and can also produce behavioral changes during recovery. Typically, we find that even at 15 minutes post-recovery from ethanol anesthesia back in its home tank, the octopus rarely takes food when offered. For these reasons, both ethanol and MgCl_2_ general anesthesia pose certain concerns for neurobehavioral studies of the intact or isolated octopus arm.

There is still debate on whether ethanol and MgCl_2_ provide “true” general anesthesia in octopuses (Abbo et al., 2021; Di Cosmo et al., 2023), particularly with regard to providing a reversible blockade of neural responses to noxious stimuli. Recent introduction of the volatile anesthetic isoflurane for octopuses, combined with pre-treatment with MgCl_2_, allows rapid recovery from insentience (Di Cosmo et al., 2023; Polese et al., 2014; Winlow et al., 2018). Isoflurane is widely used for anesthesia in mammals and fish for the same reasons. With important and appropriate safety measures, isoflurane may prove to be markedly useful for experimental procedures requiring fuller immobilization, presumed block of suffering, relatively rapid recovery, and perhaps less hangover than ethanol. However, chemical sedation of any kind in most animals is generally trailed by anhedonic affect.

Here we introduce a modified procedure of octopus tonic immobility potentially better suited than general anesthesia for certain behavioral experiments and less-invasive studies that do not involve surgery on the central nervous system (CNS).

### Tonic Immobility in Octopus

Octopus TI was first described as “hypnosis” by Vasily Danilewsky in 1890 in a description of handling-induced still responses in various animal species (Danilewsky, 1890; Zapato, 2011), and later reproduced by Jasper Ten Cate in 1928 (Ten Cate, 1928; Zapato, 2011). In such responses in our hands, octopuses assume a calm, relatively stationary state, with largely only local responses to sensory stimuli or even local MgCl_2_ injections. To our knowledge, this method has not been revisited until the present study. In the original studies, TI was induced by holding the octopus out of water for an extended time, mouth-side up and making sure that the arm suckers did not come into contact with anything to receive stimulation (Ten Cate, 1928). We have modified the hypnosis procedure to minimize stress: instead of holding the octopus out of water, it is nearly fully immersed to maintain its respiration and we use a soft net or mesh-material to gently hold the animal mouth-side up close to the surface of the water.

TI is not actually general anesthesia and cannot replace it, but in certain instances the procedure is preferable to ethanol and MgCl_2_ anesthesia, particularly for preserving local behaviors. Recovery from TI is quicker and more complete, with the animal often taking food immediately after recovery, in contrast to ethanol anesthesia. TI also avoids the sluggish and aversive behavior often following ethanol anesthesia and may also prevent potential ethanol-induced, cytotoxic tissue damage. We find that TI combined with local MgCl_2_ injection is better than ethanol anesthesia and amputation for isolating arm nerve cord from the brain and for electrode implantation. Ethanol seems to negatively affect the quality of electrophysiological activity as well as arm behavior, whereas arm amputation is followed by relatively rapid loss of sucker-arm coordinated behavior. Moreover, the TI procedure has so far never suppressed the animals’ respiration, as sometimes happens with ethanol.

While octopus TI can take more time and effort to induce than ethanol or MgCl_2_ anesthesia and appears ineffective for senescent animals, we find that this method can in some instances be a preferable alternative to the most common general anesthesia methods, that it has significant potential for promoting octopus welfare, and that it may facilitate novel neurobehavioral studies

## METHODS AND RESULTS

### Animals

Specimens of *Octopus rubescens* trapped in Monterey Bay, CA were obtained from Monterey Abalone Co. (Monterey, CA) and housed separately in artificial seawater (ASW) at 11-12°C. Specimens of *Octopus bimaculoides* were obtained from Chuck Winkler of Aquatic Research Consultants (Long Beach, CA), him, and housed separately in ASW at 18-22°C. 17 female octopus (2 *O. bimaculoides* and 15 *O. rubescens*, ranging from 4 to 200 grams) were used in these experiments. Animals were fed pieces of shrimp or squid flesh every 1-2 days. All experiments were carried out in accordance with protocol #23015 approved by the University of Illinois Urbana-Champaign (UIUC) Institutional Animal Care and Use Committee (IACUC).

### Octopus Tonic Immobility Protocol

Our modified protocol for octopus TI is demonstrated in **Supplementary Video S1** and is shown in **Figure 1**. The octopus is first placed in a large hand-net and/or soft mesh material (such as pantyhose) inside a container with aerated ASW maintained at appropriate temperature for the species (11-13°C for *O. rubescens* and 16-22°C for *O. bimaculoides*). The octopus is gently turned mouth-side up and kept close to the surface of the water by adjusting the net mesh, ensuring that the head and mantle are always submerged, and that the octopus breathes. Maintaining an optimal water level is important, such that the submersion is sufficient for breathing but not enough to fully cover their arms or mouth. If, during TI induction, the octopus attempts to right itself and move to the edge of the net, we gently move it back to the center of the net to maintain its mouth-side-up position by adjusting the net. This is repeated as needed.

**Figure 1.**
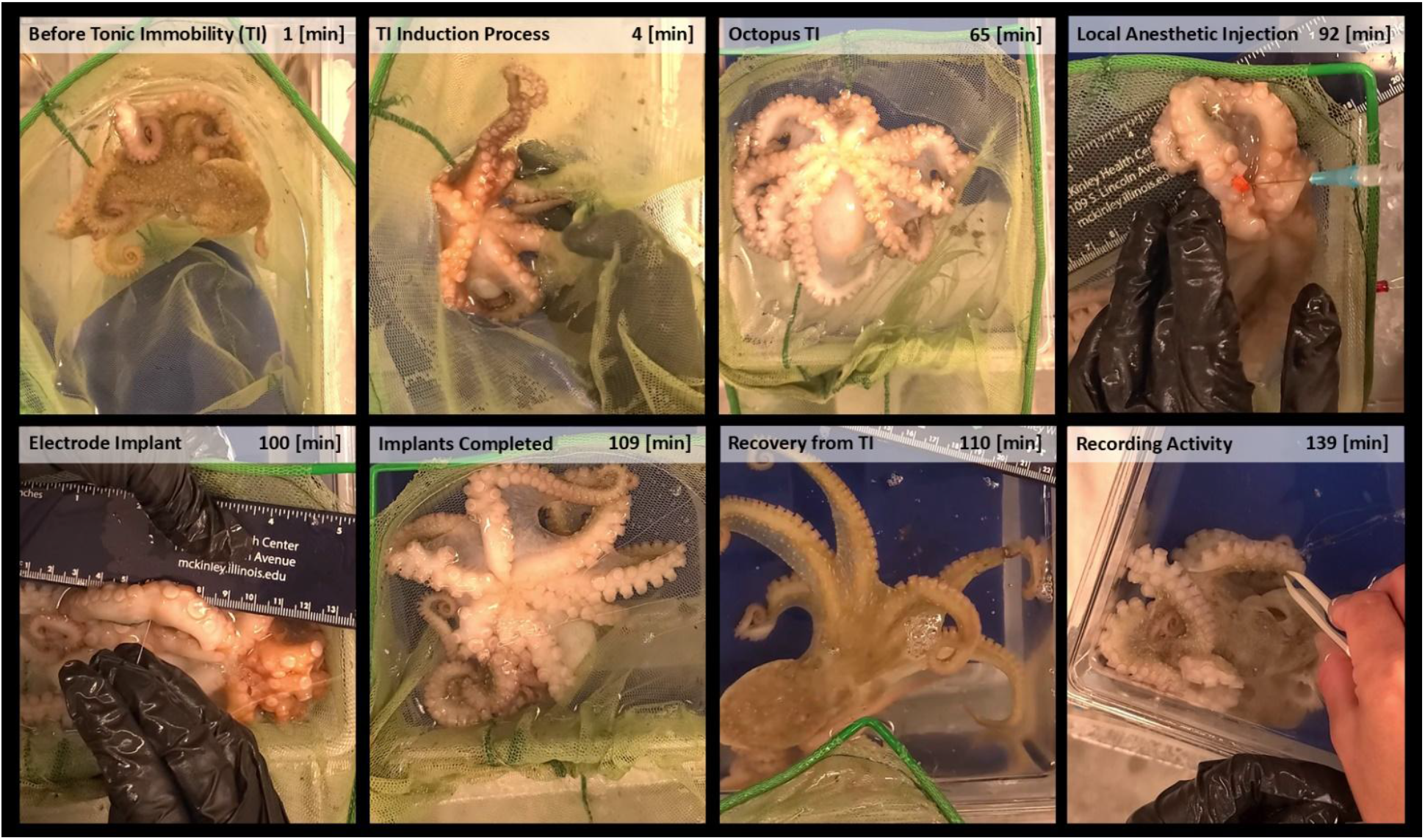
Still images from **Supplementary Video S1**, showing the octopus tonic immobility procedure and electrode implantation in an *O. rubescens*: induction of tonic immobility (t = 1 to 65 min), followed by local anesthesia with MgCl_2_ injection (t = 92 min), electrode implantation (100-109 min), recovery from tonic immobility (t = 109-110 min), and recording of neuromuscular activity from implanted electrodes (t = 110-139 min).

Typically, as TI induction continues, the octopus stays still for longer times and righting attempts are less frequent. Positioning the octopus in a cradle-like manner, keeping the arms and suckers at or above the surface of the water and unattached to surfaces, covering their eyes, and generally limiting exposure to visual stimuli appears to help TI induction. The TI induction process continues until the octopus remains mouth-side up with its arms and body still except for breathing for at least 5 minutes. Typically, for *O. rubescens*, this state is also accompanied by a marked reduction in pigmentation of the suckers and oral side of the arms. At this stage, after we observe at least 5 minutes of this still state, we test TI by lightly touching and squeezing one of the octopus’ arms. We define the TI procedure as successful if we observe only local responses in response to this arm tactile stimulation, such that only the stimulated arm moves without recruiting activity in the other arms, and the animal shows no other changes in its behavior. Notably, the responses of an immobile octopus are markedly similar to those of a sleeping octopus.

### Local Anesthesia

TI facilitates administration of local anesthetics, such as MgCl_2_ injection in an arm. Local arm anesthetization was done by injecting the base of an octopus arm by the margin of the web with iso-osmotic 330 mM MgCl_2_ solution with 10 mM HEPES buffer, pH = 7.6 (Butler-Struben et al., 2018). This appears to temporarily disrupt communication between the octopus’s arm peripheral nervous system (PNS) and its CNS. This has been tested in both immobilized and awake animals by strong tactile stimulation of the injected arm without other recruited responses in other arms or the body. Successful blockade also commonly results in a blanching of the octopus arm distal to the injection site. Typically, MgCl_2_ injection using thinner needles (30 ga), more precise depth control (aided with rubber stopper on needle) to target the arm central nerve cord, and relatively small volumes (0.1 ml for the arm sizes of the small animals we worked with here) yielded better arm anesthetization.

### Use of Octopus Tonic Immobility in Broader Experiments

#### Octopus Sucker Stimulation in TI

A marked advantage of octopus TI in behavioral studies is that local responses of the arms are preserved, unlike for general anesthesia. In a successful TI the octopus is quite still in its mouth-side-up position with suckers easily accessible and where moderate tactile stimulation produces only local arm responses that have little or no effect on all other arms or the octopus’ body. Thus, octopus TI is particularly suitable for characterizing local responses in the arm, as seen in our experiments with sucker orienting responses.

Following TI induction, sensory stimulation of suckers was done in locally anesthetized and non-anesthetized arms. A blunted syringe needle was used to precisely deliver chemotactile stimuli to octopus suckers, such as either an ASW jet or squid extract mixed with red food dye for tracking of chemical plumes. Plumes of squid extract seen to contact the arm and suckers at specific points elicited approach or avoidance responses in which the suckers extended towards or retracted away from the plume, recruited neighboring suckers, and even produced arm retraction or reaching behaviors (**Fig. 2A, left**). While a similar range of behaviors was observed in both anesthetized and non-anesthetized arms, it remains to be explored whether there are differences in approach/avoidance behaviors dependent on CNS-PNS communication, internal appetitive state of the octopus as affected by hunger and stress, as well as in distal/proximal sucker stimulation. We expect the TI procedure to facilitate further relevant experiments.

**Figure 2.**
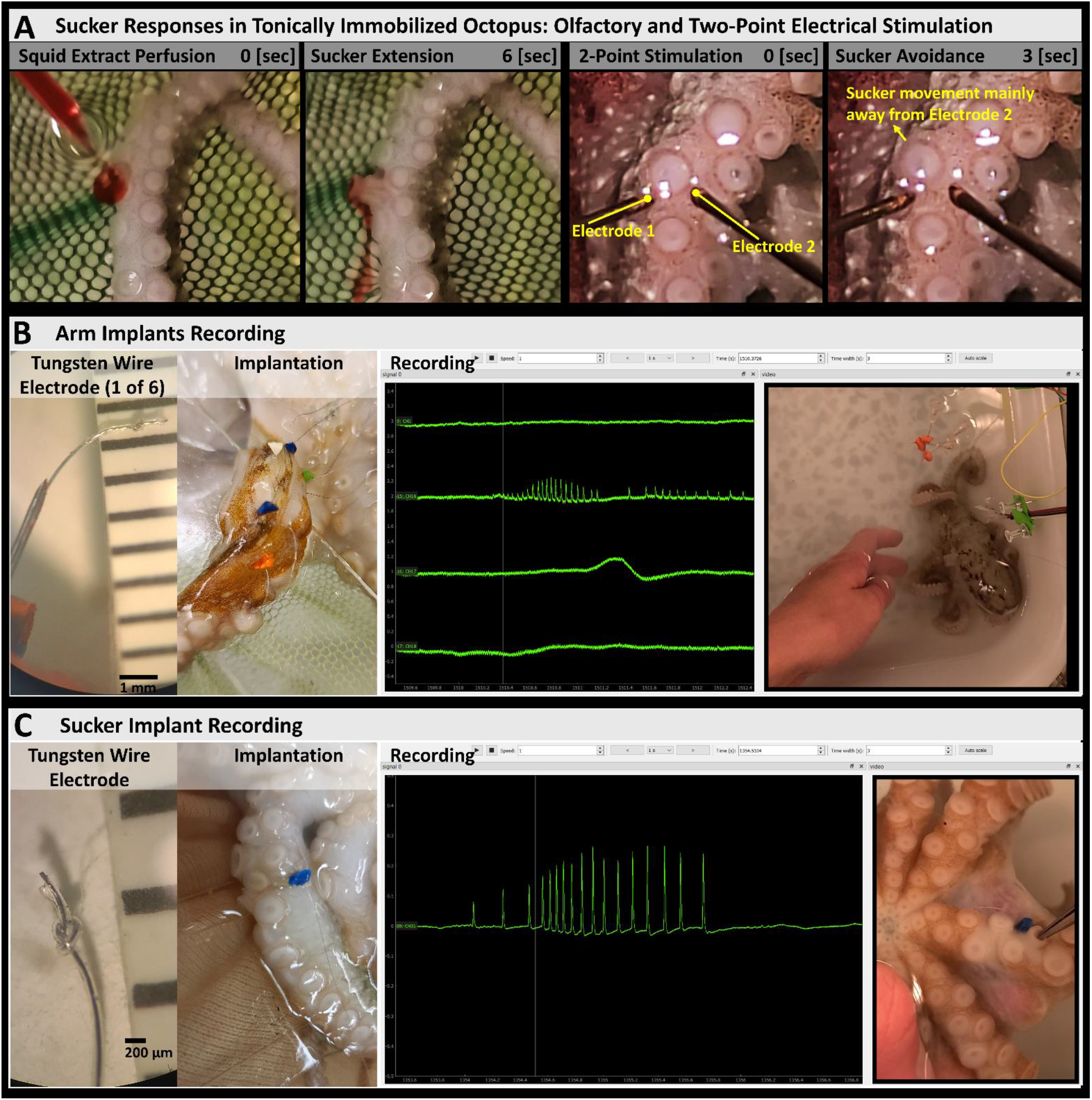
Application of octopus tonic immobility in broader experiments. **A)** Tonic immobility facilitated the testing of sucker responses to different stimuli. The left panels show a chemical stimulus (squid homogenate, with red food dye) being perfused primarily onto one sucker at t = 0 sec, followed by a precise (appetitive) extension of the sucker towards the chemical stimulus (t = 6 sec), with a few slight movements in neighboring suckers, and no noticeable responses in other arms. The right panels show two bipolar concentric electrodes positioned on the outer rim of a sucker (labels in yellow). A train of electrical pulses (at 10 Hz, each pulse 4 V and 1.5 ms duration) was delivered through both electrodes simultaneously starting at t = 0 sec and lasting for about 1.5 sec. This caused aversive movement of the sucker (t = 3 sec), directly away from the rightmost electrode. **B)** Octopus tonic immobility was combined with local anesthesia (MgCl_2_ injection) to implant tungsten wire electrodes in the octopus arm (insertion points labeled with colored rubber markers) and obtain electrophysiological recordings in a freely-moving octopus. **C)** Tonic immobility facilitated implantation of a tungsten wire electrode in an octopus sucker (insertion point labeled with a blue rubber markers), as well as for testing sucker electrophysiological responses with tactile stimulation.

Previously we and others have observed that responses in arms isolated by amputation decline rapidly, impeding tests of sucker responses to chemotactile stimulation. Therefore, we tested arms under TI. MgCl_2_ was injected under TI in a proximal arm site to isolate it from the CNS. Keeping the octopus in a convenient position for extended time periods, we conducted experiments with one- and two-point stimulation with bipolar concentric electrodes and a Grass S-5 stimulator to deliver shocks (bipolar pulses of 1.5 ms duration, ranging 1-6 V in amplitude, in single-pulse regime or 10 Hz train of pulses ranging in duration from 0.5-3 sec) at one or two different sites on a single sucker’s outer rim (**Fig. 2A, right**). A sucker responds to a single point stimulus (tactile, chemical, or electrical) on its outer rim with a very precise movement directly towards or away from the stimulus. It appeared that the sucker could estimate the bearing to a single point stimulus and use it to generate very accurate motor output for approach/avoidance. We tested arms in intact animals for two-point stimulation to see whether suckers might average the stimuli loci in their directional responses. While data are still being gathered, early results suggest suckers produce a winner-take-all response (Fukai & Tanaka, 1997; Maass, 2000) to only one of the two loci stimulated simultaneously, rather than averaging the two response sites to move at an angle between them as has been observed in turn responses of the sea slug *Pleurobranchaea californica* (Yafremava et al., 2007; Yafremava & Gillette, 2011).

These results are consistent with neuronal circuit architectures in the sucker like lateral inhibition (Mao & Massaquoi, 2007; von Békésy, 1967) and are useful in modeling work. In future, we hope to elucidate the mechanisms behind sensory processing in the local circuitry of the sucker, as well as recruitment of neighboring suckers.

### Electrophysiology in Freely-Moving Octopuses

Tungsten wire electrodes were implanted into octopus arms to record electrical activity. Electrodes were PFA-coated tungsten wire (0.1 mm diameter, A-M Systems) with a terminal half-hitch knot to anchor (**Fig. 2B-C**). For implantation (**Supplementary Video S1, Fig. 1**) after TI induction, chilled 330 mM MgCl_2_ solution was injected at the base of the arm to disrupt arm-brain communication. The wire electrodes were carefully inserted into the tissue coaxially within a 30 ga needle (with an optional rubber stopper on the needle for additional control over implantation depth) (Gutfreund et al., 1998; Parker, 1968; Uyeno & Kier, 2007). Data were recorded with a headstage (Intan RHD 32-channel) and an acquisition board (Open Ephys) with a sampling rate of 20 kS/s and amplifier hardware bandwidth from 0.1 to 7600 Hz. Videos were recorded with a phone camera at 30 fps, and synchronized with electrophysiological recordings using a simultaneous visual and electrical signal: the flashing of a light-emitting diode (LED) coupled with a 5 V stimulator. Synchronized recordings were visualized and reviewed using the Python library ephyviewer (Garcia & Gill, 2021; Gill et al., 2020) (Ephyviewer ver. 1.5.1, Python ver. 3.9.7), with the stand-alone viewer slightly modified to display both electrophysiological recordings and video. Modified code for viewing synchronized recordings and video is available at https://github.com/KatyaGribkova/ephyviewer.

The arm implant recordings showed spiking activity during arm movements in a freely moving octopus, while an implant at the base of the sucker recorded the most activity with inner-central tactile stimulation of the sucker (**Fig. 2B-C**). We expect the recordings in both cases to represent compound action potentials of muscle fibers, similar to those observed in the electrode implants of Gutfreund et al. (1998). To our knowledge, this may be a first successful sucker electrode implantation and recording. Previously, we had encountered difficulties in obtaining electrophysiological recordings from tungsten wire electrodes implanted under 1% ethanol anesthesia (see notes in **Supplementary Data S1**, 11/14/2022), which was one of the main motivations to implement the tonic immobility procedure, alongside local anesthesia, in the implantation process.

### Variability in Tonic Immobility Onset and Duration

There was appreciable variability in latency to TI onset as well as TI duration across the animals we worked with. We define “latency to TI onset” as the time taken for the octopus to achieve an uninterrupted still state of at least 5 minutes, and with only local responses to stimuli as described earlier in the Octopus Tonic Immobility Protocol. “TI duration” is the length of time that TI was maintained in any particular experiment, whether spontaneously recovered or interrupted by the experimenter to test procedures (electrode implantation, sucker stimulation, etc.).

Multiple factors may affect latency of TI onset, including octopus species, age, size, circadian rhythm, senescence, hunger, and other characteristics and dynamic internal states of the octopus. Notably, We observed a significant positive correlation between octopus weight and latency of TI onset (**Fig. 3A**, Spearman’s rank correlation: r_S_ = 0.600, p < 0.002). This is explainable in a number of ways, including difficulty in handling larger octopuses as well as secondary factors that also correlate with size, like age and senescence.

**Figure 3.**
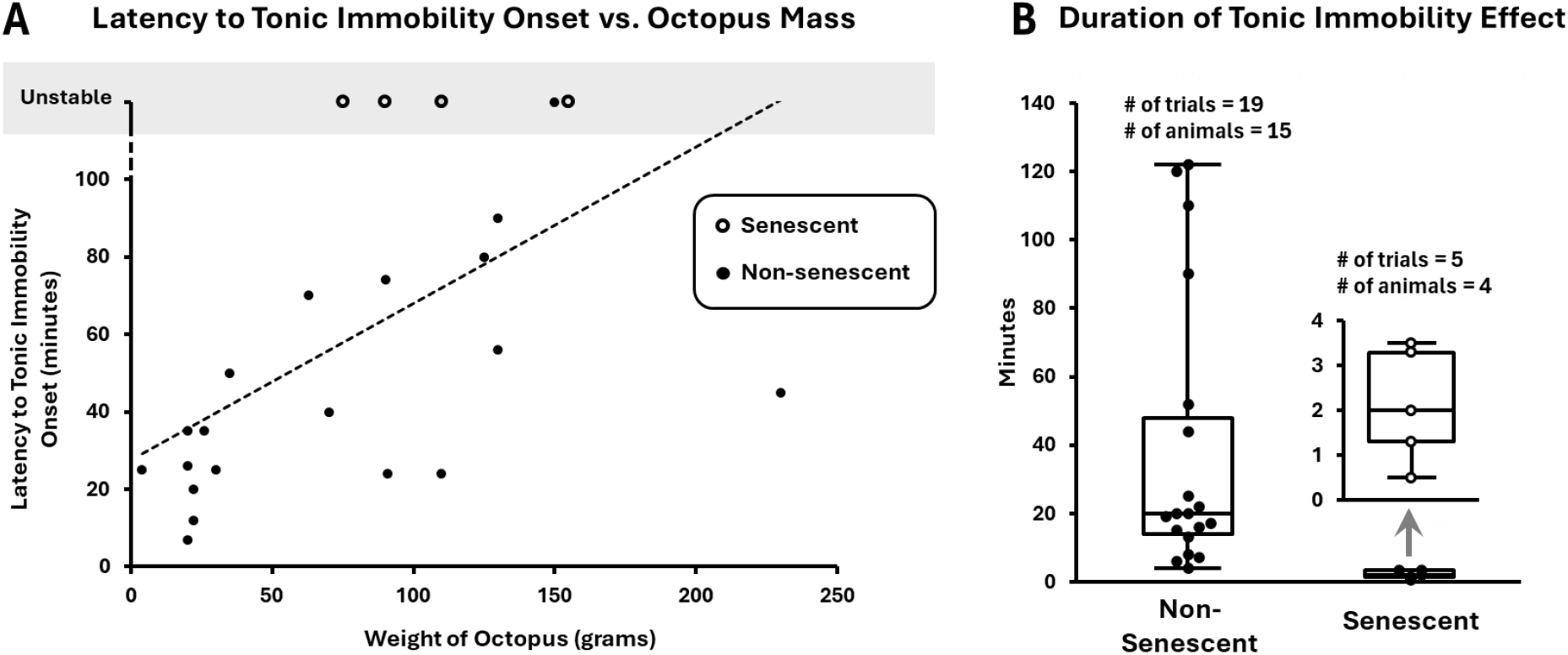
Exploring the variability of octopus tonic immobility onset and duration. **A)** Latency to tonic immobility onset has a strong positive correlation with approximate octopus mass (Spearman’s rank correlation: r_S_ = 0.600, p = 0.00194). **B)** While the number of observations for clearly senescent octopuses is presently small, tonic immobility for these animals was quite unstable: still states were typically maintained for less than 5 minutes (mean 2.12 min, std. dev. 1.15) whereas in almost all non-senescent animals still states were maintained for at least 5 minutes (mean 38.4 min, std. dev. 39.4), and up to 2 hrs, with a significant difference (2-tailed Mann-Whitney U test: U = 1, z-score = 3.34, p = 0.00084). The inset shows a closer look at the box-and-whisker plot of tonic immobility effect duration for senescent octopuses.

In particular, while our number of observations for clearly senescent octopuses (Anderson et al., 2002; Hanlon, 1983; Roumbedakis & Guerra, 2019; Roura et al., 2024; Wodinsky, 1977) (1 *O. bimaculoides*, and 3 *O. rubescens*, all of which had stopped taking food after having laid eggs), was small (N=4), we observed striking differences when attempting to induce TI with these animals as compared to non-senescent animals (Mann-Whitney U test for duration of TI effect: U = 0, p < 0.001). In 5 trials with these senescent octopuses, we were unable to maintain a still state for more than 5 minutes (**Fig. 3B**), even though most of our attempts lasted more than 2 hrs (**Fig. 3A, unstable**). Notably, in one of these animals, a large female *O. rubescens*, when the octopus was not yet senescent, a still state was successfully maintained for at least 40 minutes in 3 different trials. But once this animal had laid eggs, turned senescent, and was no longer taking food, we were unable to maintain a still state for more than 4 minutes. In another curious case, we observed a female *O. rubescens* that had laid sterile eggs, but was not paying particular attention to them and was still taking food. At this stage, we were able to induce a stable still state with her after about 24 minutes, that lasted for at least 22 minutes. However, about 17 days later, this octopus had stopped taking food, and we were then unable to induce TI with her for more than 4 minutes, even though the attempt lasted for more than 2 hrs. Detailed observations of all TI procedures are included in **Supplementary Data S1**.

While senescent animals could be sufficiently calmed to conduct certain experiments, such as testing of sucker orienting responses, they seemed much more likely than non-senescent animals to respond to light arm touches or squeezes with more general responses involving recruitment of other arms. This could be due to sensorimotor and neurobehavioral changes that occur with onset of senescence in octopuses, particularly those associated with the care and guarding of octopus eggs (Anderson et al., 2002; Roumbedakis & Guerra, 2019; Wang & Ragsdale, 2018). The octopus optic gland plays an important role in regulating these reproductive behaviors, with removal of optic glands in senescent octopus often resulting in the animal taking food again and delaying senescence-related starvation (Anderson et al., 2002; Wang & Ragsdale, 2018; Wodinsky, 1977). In particular, we note the successful TI with the *O. rubescens* that still took food after laying eggs, which was then followed by unsuccessful TI attempts when this octopus had stopped taking food. This case raises some interesting questions about how optic gland signaling might affect octopus sensorimotor processing and behaviors during TI induction, in addition to feeding behaviors, and how it affects the efficacy of the TI procedure. Further, octopuses in decline also seem to lose their vestibulo-ocular reflex (Wang & Ragsdale, 2018) that would normally orient their pupils horizontally with respect to gravity, which may also have an effect on TI induction.

## DISCUSSION

Tonic immobility is notable for its inducibility in many animals, including humans, through restraint or, in some cases, threat, and often involves mild analgesia (Carli, 1978; Carli & Farabollini, 2022a, 2022b). The struggles of a potential prey can intensify an attacker’s effort (Gillette et al., 2000), and TI is generally characterized as a defensive response where inactivity may reduce a predator’s interest. It may also reduce further stress by shutting down sensory input to CNS.

While Danilewsky’s (1890) original report described the still reaction of the octopus as hypnosis, the hypnotic state in humans is induced differently by voluntary concentration and represents a focused state of attention. The early investigator James Braid (Braid, 1845) characterized the human state as “…the induction of a habit of intense abstraction, or concentration of attention…). More recent human fMRI studies of the cortical networks for ideation vs. attention show that hypnosis is associated with lesser activity in the default mode network and that high hypnotizability is associated with greater functional connectivity between the (attentional) executive control network and the salience network, and that: “These changes in neural activity underlie the focused attention, enhanced somatic and emotional control, and lack of self-consciousness that characterizes hypnosis.” (Jiang et al., 2017). Thus, the octopus state of tonic immobility does not admit the voluntary behavior and responsivity of the human hypnotic state.

In octopuses, induced TI as we describe has potential advantages over general anesthesia in certain cases, particularly in experiments where preservation of local arm behavior is needed, and where strongly invasive surgical procedures are not necessary. Handling unaccustomed individuals of many species is typically stressful and general anesthesia may compound that. However, the very rapid recovery of the octopus from TI on righting and approach to food contrasts with the slower recoveries from other anesthetic methods.

While general anesthesia with ethanol and MgCl_2_ should remain useful in octopus research until better ways are found, drawbacks of these methods can be avoided with TI in certain experiments. Notably, general anesthesia affects both CNS and PNS function and impedes testing of local sensorimotor responses. To our knowledge, our modification of octopus TI (**Fig. 1**) may be the only reported method to sufficiently calm non-senescent octopus to test local sensorimotor responses. The induced TI appears similar to the tonic immobility seen in various other species (Carli & Farabollini, 2022b, 2022a), such as sharks (Páez et al., 2023), although unlike probably most of these cases, octopus TI does not significantly suppress some of the local peripheral responses. With TI, even without local anesthesia, there seems to be decreased communication between octopus CNS and PNS, such that lightly squeezing the arm of a sufficiently torpid octopus generally produces only local responses, without responses in other arms or body. This effect facilitates local injection of anesthetic MgCl_2_ solution at the base of arms, and since the octopus lies still in its mouth-side-up position, it offers a markedly convenient opportunity for testing local arm and sucker responses (**Fig. 2A**). Thus, isolation of the arm from the brain and the rest of the PNS to test autonomous behavior may be easily gotten without amputation in TI animals by local MgCl_2_ injection. Further, the combination of local anesthesia and TI greatly facilitates implantation of electrodes for recording neuromuscular activity (**Fig. 1, Fig. 2B-C**). While we currently have a very limited number of two-point stimulation and arm implant recordings, further studies would be quite useful for understanding CNS-PNS communication and sensory processing in TI, and for comparing it against methods of anesthesia. Lastly, in our various experiments, we noted interesting variability in octopus TI onset and duration, closely correlated with both octopus size and senescence (**Fig. 3**). TI induction may basically be ineffective for senescent animals; however, further testing may usefully explore this difference.

The modified octopus TI procedure may therefore provide new opportunities to advance both octopus laboratory welfare and neurobehavioral studies. There yet remains much to be learned about the mechanisms that give rise to the still reaction, what particular neurophysiological effects it may induce in CNS-PNS communication, and if there are any analgesic effects similar to those observed in TI with other species (Carli, 1978; Carli & Farabollini, 2022b, 2022a). As the procedure involves holding the animals mouth-side up, and since it seems ineffective for senescent animals which may have lost their vestibulo-ocular reflex (Wang & Ragsdale, 2018), the vestibular system might be involved in TI induction. Whatever the mechanisms, if ways could be found to manipulate them directly it might further facilitate experiments to further unravel details of the complex brains and behaviors of these intriguing creatures.

## ACKNOWLEDGEMENTS

We are grateful to Dr. William Gilly for hosting us at Hopkins Marine Station where we performed our first octopus TI experiments, and to Trevor Fay, Art Seavey, Caitlin Rooney and the other staff of Monterey Abalone Co., and Chuck Winkler of Aquatic Research Consultants for their efforts in animal supply. We also thank our dedicated undergraduate students, including Emily Stine, Tyler T. Cushing, Trisha Pal, Carina B. Dudick, Paige Henley for learning the TI method and using it in their experiments.

## AUTHOR CONTRIBUTIONS

Ekaterina D. Gribkova: Conceptualization, Formal analysis, Investigation, Methodology, Software, Validation, Visualization, Writing – original draft, Writing – review & editing. Jilai Cui: Investigation, Methodology, Writing – review & editing. Rhanor Gillette: Conceptualization, Funding acquisition, Investigation, Methodology, Resources, Visualization, Writing – original draft, Writing – review & editing.

## DATA AVAILABILITY

The Python library ephyviewer (Garcia & Gill, 2021; Gill et al., 2020) (Ephyviewer ver. 1.5.1, Python ver. 3.9.7), was modified for additional data visualization. The modified code is available at https://github.com/KatyaGribkova/ephyviewer.

## ETHICAL CARE CONSIDERATIONS

Specimens of *Octopus rubescens* trapped in Monterey Bay, CA were obtained from Monterey Abalone Co. (Monterey, CA) and housed separately in artificial seawater (ASW) at 11-12°C. Specimens of *Octopus bimaculoides* were obtained from Chuck Winkler of Aquatic Research Consultants (Long Beach, CA), and housed separately in ASW at 18-22°C. 17 female octopus (2 *O. bimaculoides* and 15 *O. rubescens*, ranging from 4 to 200 grams) were used in these experiments. Animals were fed pieces of shrimp or squid flesh every 1-2 days. All experiments were carried out in accordance with protocol #23015 approved by the University of Illinois Urbana-Champaign (UIUC) Institutional Animal Care and Use Committee (IACUC).

### Legend for Supplementary Video 1

This video shows our modified procedure for octopus “hypnosis”, or tonic immobility, its use in electrode implantation in octopus arms, and lastly the octopus’s quick recovery from tonic immobility.

## REFERENCES

Abbo, L. A., Himebaugh, N. E., DeMelo, L. M., Hanlon, R. T., & Crook, R. J. (2021). Anesthetic efficacy of magnesium chloride and ethyl alcohol in temperate octopus and cuttlefish species. In Journal of the American Association for Laboratory Animal Science (Vol. 60, Issue 5, pp. 556–567).

Anderson, R. C., Wood, J. B., & Byrne, R. A. (2002). Octopus senescence: The beginning of the end. In Journal of Applied Animal Welfare Science (Vol. 5, Issue 4, pp. 275–283).

Andrews, P. L. R., & Tansey, E. M. (1981). The effects of some anaesthetic agents in Octopus vulgaris. Comparative Biochemistry and Physiology Part C: Comparative Pharmacology, 70(2), 241–247. 10.1016/0306-4492(81)90057-5

Braid, J. (1845). Mr. Braid on Hypnotism. The Lancet, 45(1135), 627–628. 10.1016/S0140-6736(02)65024-X

Butler-Struben, H. M., Brophy, S. M., Johnson, N. A., & Crook, R. J. (2018). In vivo recording of neural and behavioral correlates of anesthesia induction, reversal, and euthanasia in cephalopod molluscs. In Frontiers in physiology (Vol. 9, p. 109).

Carli, G. (1978). Animal Hypnosis and Pain. In F. H. Frankel & H. S. Zamansky (Eds.), Hypnosis at its Bicentennial (pp. 69–77). Springer US. 10.1007/978-1-4613-2859-9_6

Carli, G., & Farabollini, F. (2022a). Defensive responses in invertebrates: Evolutionary and neural aspects. In Progress in Brain Research (Vol. 271, Issue 1, pp. 1–35).

Carli, G., & Farabollini, F. (2022b). Pain control in tonic immobility (TI) and other immobility models. Progress in Brain Research, 271(1), 253–303.

Crook, R. J., Hanlon, R. T., & Walters, E. T. (2013). Squid have nociceptors that display widespread long-term sensitization and spontaneous activity after bodily injury. In Journal of Neuroscience (Vol. 33, Issue 24, pp. 10021–10026).

Danilewsky, B. (1890). Recherches physiologiques sur l’hypnotisme des animaux. In Congrès international de psychologie. Compte rendu des séances. P (pp. 79–92).

Di Cosmo, A., Maselli, V., Cirillo, E., Norcia, M., de Zoysa, H. K., Polese, G., & Winlow, W. (2023). The Use of Isoflurane and Adjunctive Magnesium Chloride Provides Fast, Effective Anaesthetization of Octopus vulgaris. In Animals (Vol. 13, Issue 22, p. 3579).

Fukai, T., & Tanaka, S. (1997). A simple neural network exhibiting selective activation of neuronal ensembles: From winner-take-all to winners-share-all. In Neural computation (Vol. 9, Issue 1, pp. 77–97).

Garcia, S., & Gill, J. P. (2021). Ephyviewer [Python].

Gill, J. P., Garcia, S., Ting, L. H., Wu, M., & Chiel, H. J. (2020). neurotic: Neuroscience tool for interactive characterization. In Eneuro (Vol. 7, Issue 3).

Gillette, R., Huang, R.-C., Hatcher, N., & Moroz, L. L. (2000). Cost-benefit analysis potential in feeding behavior of a predatory snail by integration of hunger, taste, and pain. Proceedings of the National Academy of Sciences, 97(7), 3585–3590. 10.1073/pnas.97.7.3585

Grimm, K. A., Lamont, L. A., Tranquilli, W. J., Greene, S. A., & Robertson, S. A. (2015). Veterinary anesthesia and analgesia. John Wiley & Sons.

Gutfreund, Y., Flash, T., Fiorito, G., & Hochner, B. (1998). Patterns of arm muscle activation involved in octopus reaching movements. In Journal of Neuroscience (Vol. 18, Issue 15, pp. 5976–5987).

Hanlon, R. (1983). Octopus joubini. In Cephalopod life cycles (Vol. 1, pp. 293–310).

Jiang, H., White, M. P., Greicius, M. D., Waelde, L. C., & Spiegel, D. (2017). Brain activity and functional connectivity associated with hypnosis. Cerebral Cortex, 27(8), 4083–4093.

Maass, W. (2000). On the computational power of winner-take-all. In Neural computation (Vol. 12, Issue 11, pp. 2519–2535).

Mao, Z.-H., & Massaquoi, S. G. (2007). Dynamics of winner-take-all competition in recurrent neural networks with lateral inhibition. IEEE Transactions on Neural Networks, 18(1), 55–69.

Páez, A. M., Padilla, E. M. H., & Klimley, A. P. (2023). A review of tonic immobility as an adaptive behavior in sharks. Environmental Biology of Fishes, 106(6), 1455–1462.

Parker, T. G. (1968). Simple method for preparing and implanting fine wire electrodes. In American Journal of Physical Medicine & Rehabilitation (Vol. 47, Issue 5, pp. 247–249).

Perez, P. V., Butler-Struben, H. M., & Crook, R. J. (2017). The selective serotonin reuptake inhibitor fluoxetine increases spontaneous afferent firing, but not mechanonociceptive sensitization, in octopus. In Invertebrate Neuroscience (Vol. 17, pp. 1–5).

Polese, G., Winlow, W., & Di Cosmo, A. (2014). Dose-dependent effects of the clinical anesthetic isoflurane on Octopus vulgaris: A contribution to cephalopod welfare. In Journal of aquatic animal health (Vol. 26, Issue 4, pp. 285–294).

Roumbedakis, K., & Guerra, Á. (2019). Cephalopod senescence and parasitology. In Handbook of pathogens and diseases in cephalopods (pp. 207–211).

Roura, Á., Castro-Bugallo, A., Martínez-Pérez, M., & Guerra, Á. (2024). Senescence in common octopus, Octopus vulgaris: Morphological, behavioural and functional observations. In Applied Animal Behaviour Science (p. 106294).

Ten Cate, J. (1928). Nouvelles observations sur l’hypnose dite animale. Etat d’hypnose chez octopus vulgaris. In Archives néerlandaises de physiologie de l’homme et des animaux (Vol. 13, pp. 402–406).

Uyeno, T. A., & Kier, W. M. (2007). Electromyography of the buccal musculature of octopus (Octopus bimaculoides): A test of the function of the muscle articulation in support and movement. In Journal of Experimental Biology (Vol. 210, Issue 1, pp. 118–128).

von Békésy, G. (1967). Mach band type lateral inhibition in different sense organs. In The Journal of general physiology (Vol. 50, Issue 3, pp. 519–532).

Wang, Z. Y., & Ragsdale, C. W. (2018). Multiple optic gland signaling pathways implicated in octopus maternal behaviors and death. Journal of Experimental Biology, 221(19), jeb185751.

Winlow, W., Polese, G., Moghadam, H.-F., Ahmed, I. A., & Di Cosmo, A. (2018). Sense and insensibility–an appraisal of the effects of clinical anesthetics on gastropod and cephalopod molluscs as a step to improved welfare of cephalopods. Frontiers in Physiology, 9, 1147.

Wodinsky, J. (1977). Hormonal inhibition of feeding and death in octopus: Control by optic gland secretion. In Science (Vol. 198, Issue 4320, pp. 948–951).

Yafremava, L. S., Anthony, C. W., Lane, L., Campbell, J. K., & Gillette, R. (2007). Orienting and avoidance turning are precisely computed by the predatory sea-slug Pleurobranchaea californica McFarland. Journal of Experimental Biology, 210(4), 561–569.

Yafremava, L. S., & Gillette, R. (2011). Putative lateral inhibition in sensory processing for directional turns. Journal of Neurophysiology, 105(6), 2885–2890. 10.1152/jn.00124.2011

Zapato, L. (2011, June 5). How To Hypnotize An Octopus [Blog]. ZPi. https://zapatopi.net/blog/?post=201105068839.how_to_hypnotize_an_octopus

